# Exploring links between climatic predictability and the evolution of within- and transgenerational plasticity

**DOI:** 10.1101/2022.03.20.484890

**Authors:** Sridhar Halali, Marjo Saastamoinen

## Abstract

In variable environments, phenotypic plasticity can increase fitness by providing tight environment-phenotype matching. However, adaptive plasticity is expected to evolve only when the future selective environment can be predicted based on the prevailing conditions. That is, the juvenile environment should be predictive of the adult environment (within-generation plasticity) or the parental environment should be predictive of the offspring environment (transgenerational plasticity). Here, we test links between environmental predictability and evolution of adaptive plasticity by combining time series analyses and a common garden experiment using temperature as a stressor in a temperate butterfly (*Melitaea cinxia*). Time series analyses revealed that across season fluctuations in temperature over 48 years is overall predictable. However, within the growing season, temperature fluctuations showed high heterogeneity across years with low autocorrelations and timing of temperature peaks were asynchronous. Most life-history traits showed strong within-generation plasticity for temperature and traits such as body size and growth rate broke the temperature-size rule. Evidence for transgenerational plasticity, however, was weak and detected for only two traits each in an adaptive and non-adaptive direction. We suggest that low predictability of temperature fluctuations within the growing season likely disfavours the evolution of adaptive transgenerational plasticity but instead favours strong within-generation plasticity.

## INTRODUCTION

Phenotypic variation in the wild can be shaped by genetics or by the environment, but often it is an interaction (G × E) of both of these factors. Traditionally, evolutionary biologists have had a strong gene-centric view in adaptive evolution (Pigliucci 2005, Bonduriansky 2021) but the field has witnessed a renaissance in recent years with respect to the contribution of non-genetic factors at both micro- and macroevolutionary scales (Bonduriansky 2012, Pfennig et al. 2010, Levis and Pfennig 2016, Jablonka 2017). One such important non-genetic phenomenon is transgenerational plasticity (henceforth TGP), where conditions experienced by parents can shape offspring fitness in both adaptive and non-adaptive manner (Mousseau and Fox 1998, Uller 2008, Bonduriansky 2021). The occurrence of TGP has been widely documented across plants and animals including humans (Uller et al. 2013, Yin et al. 2019), although its adaptive potential has been convincingly demonstrated in relatively few studies (see Galloway and Etterson 2007, Fox et al. 1997, Agrawal et al. 1999). Detecting adaptive TGP can be difficult and is hindered by factors such as lack of information on species ecology, not using ecologically relevant stressors in the experiments, and meagre information on how stressor(s) of interest affects fitness in the wild.

Most organisms live in temporally fluctuating environments, although the degree of fluctuations can differ dramatically across habitats, geographic locations and so on. When such temporal fluctuations are cyclical, for example alternating wet-dry seasons in the tropics or summer-winters in temperate regions, organisms can make use of the environmental cues (e.g. temperature or photoperiod) to predict the forthcoming selective environment (Reed et al. 2010, Chevin and Lande 2015). In such predictably fluctuating environments, phenotypic plasticity or within-generation plasticity where cue(s) experienced during juvenile development can trigger alternative developmental pathways, can be adaptive by providing a tight environment-phenotype matching (Moran 1992, Beldade et al. 2011, Bonamour et al. 2019). Seasonal polyphenism in butterflies is a classic example of such adaptive within-generation plasticity (Shapiro 1976). Apart from such clear-cut examples, plasticity can also comprise of rapid behavioral and physiological adjustments, for example, as a response to stressful conditions. Even for such transient responses, theory suggests that selection can fine-tune stress responses when stress-inducing episodes are predictable (Taborsky et al. 2021).

As the predictability of temporal fluctuations is a pre-requisite for the evolution of adaptive within-generation plasticity, similarly, adaptive TGP is expected to evolve when conditions experienced by parents are predictive of the environment experienced by their offspring (Uller 2008, Burgess and Marshall 2014, Leimar and McNamara 2015, Bonduriansky 2021). In this way, adaptive TGP can provide a jump-start in enhancing offspring’s fitness by shaping their phenotype much earlier during the development (Bell and Hellman 2019). Such type of TGP is equivalent to the anticipatory plasticity than mere condition transfer from parents to offspring (Burgess and Marshall 2014). For example, in *Caenorhabditis elegans*, the evolution of adaptive maternal effect (maternal glycogen provisioning to embryos in anoxic conditions) only occurred when normal and oxygen deprivation conditions fluctuated predictably (Dey et al. 2016, also see Lind et al. 2020). Moreover, when conditions experienced by parents are not predictive of the offspring’s environment, TGP can be even maladaptive, and selection is instead expected to favour within-generation plasticity which may allow a more immediate response to prevailing environmental conditions (Leimar and McNamara 2015, Bonduriansky 2021). Despite environmental predictability being a core pre-requisite in the evolution of both adaptive within-generation plasticity and TGP (Moran 1992, Reed et al. 2010, Leimar and McNamara 2015), thorough analysis of predictability of a relevant stressor(s) using long-term data remains scant (but see Colicchio and Herman 2020, Halali et al. 2021). It is even suggested that failure in quantifying the extent of predictability between parent-offspring environments has hindered our understanding of the evolution of adaptive TGP (Uller et al. 2013, Burgess and Marshall 2014).

Here, by using a combination of environmental time series analyses and a common garden experiment, we investigate how the predictability of the temperature (a stressor in our study) drives within-generation plasticity and TGP using the Glanville Fritillary butterfly (*Melitaea cinxia*) from the Åland archipelago (south-west Finland) as a model system. Temperature is an ecologically important stressor and a selective factor that can strongly affect life-history traits (Atkinson 1994, Kingsolver and Huey 2007). It is also a highly relevant and most explored environmental factor in the current scenario of global change (e.g. Ma et al. 2021). Moreover, microclimatic variation in temperature has been shown to have strong effects on key life-history traits in *M. cinxia* in both field and laboratory settings (Verspagen et al. 2020, Rytteri et al. 2021). Here, we use 48 years (1972-2020) of data on daily temperatures to measure the predictability of temperature fluctuations across and within growing seasons spanning parent and offspring growth periods. We then carry out a common garden experiment to investigate the extent of within-generation plasticity and the prevalence of adaptive TGP. We expect that treatments with matching parent-offspring temperatures would have higher performance (e.g. higher growth rates) compared to treatments with unmatching temperatures under the evolution of adaptive TGP (for similar findings see Salinas and Munch (2012) and Zizzari and Ellers (2014)). Overall, linking environmental predictability to the results from the experiment allows us to rigorously test the core prediction from the life-history theory that adaptive TGP is expected to evolve when parental conditions can predict environmental conditions experienced by offspring.

## METHODOLOGY

### Study species

*Melitaea cinxia* in Åland islands inhabits a large network of fragmented dry meadows (called patches) and is one of the classic model systems for metapopulation research (reviewed in Ovaskainen and Saastamoinen 2018). Finnish *M. cinxia* is univoltine and the life cycle is as follows. Adults emerge in June and females lay eggs in batches on two commonly used hostplants *Plantago lanceolata* and *Veronica spicata* (Wahlberg 2000). Pre-diapause larvae (1^st^ to 4/5^th^ instar) hatch in late June/early July and feed gregariously. In late August, larvae spin a communal web (or winter nest) usually comprising of full-sibs in which they diapause until spring. Post-diapause larvae (4/5^th^ to 7^th^ instar) recommence their development in early April and disperse to nearby hostplants. Final instar larvae and pupae in the field are typically found in May (Rytteri et al. 2021).

### Quantifying the predictability of temperature fluctuations

The predictability of temperature fluctuations across seasons (i.e. rise and fall of temperature across seasons) and within a growing season was quantified using three different approaches. Mean daily air temperature (the average temperature based usually on 4-8 observations per day) available from January 1972 to December 2020 were downloaded from the meteorological station (location:Åland islands, Jomalaby, coordinates: 60.17824, 19.98686; https://en.ilmatieteenlaitos.fi/download-observations#!/). Temperature values were missing for 22 days across the time series and these gaps were filled with approximated values using ‘*na*.*approx’* function from the *Zoo* ver 1.8.9 R package (Zeileis and Grothendieck 2005). All analyses were performed in R ver 4.1.1 (R Core Team 2021). R packages *tidyverse* ver 1.3.1 (Wickham et al. 2019) and *ggplot2* ver 3.3.5 (Wickham 2016) were used for general data handling and producing figures, respectively.

#### Wavelet analysis

Wavelet analysis or multi-resolution analysis is a powerful approach that allows quantifying any seasonal phenomenon (Cazelles et al. 2008, Tonkin et al. 2017, Rösch and Schmidbauer 2018a). Wavelet analysis offers an elegant way to capture both low and (transient) high-frequency signals in the time series by simultaneously dilating, contracting, and adjusting the height of the mother wavelet (Cazelles et al. 2008, Cazelles et al. 2014). The dilated wavelet better captures lower frequencies, while the contracted wavelet (simultaneously adjusting its height) better captures high-frequency signals. Wavelet analysis, therefore, is well suited for detecting abrupt changes in the frequencies such as sharp rise or fall of temperature in the time series (Cazelles et al. 2008, Burgess and Marshall 2014, Tonkin et al. 2017).

Wavelet analysis was carried out using the *WaveletComp* ver 1.1 (Rösch and Schmidbauer 2018b) for daily mean temperature time series from 1972 to 2020. The periodicity of the temperature fluctuation across years was visualised using wavelet spectrum (function: *wt*.*image*) and the significant periods (p<0.05 which were determined based on 100 simulations) are indicated by white ridges. Warm colours within the white ridges indicate the regions of the highest power, which represents the magnitude of variance at a given wavelet scale or simply the regions where dominant frequencies oscillate (Rösch and Schmidbauer 2018a). Moreover, average wavelet power (function: *wt*.*avg*) was plotted to determine which period of the time series showed the highest power. In addition to the temperature, wavelet analysis was performed on the photoperiod, a perfectly predictable cue, for the above specified years. This allows comparing the extent of predictability in both variables. Time series data for photoperiod for the same location was obtained using the *maptools* package ver (Bivand and Lewin-Koh 2021).

#### Temporal autocorrelations and Symbolic Aggregation Approximation algorithm

Predictability of temperature within the growing period (April to August) was measured by quantifying temporal autocorrelations and using the Symbolic Aggregation Approximation (SAX) algorithm. Post-diapause larvae usually start growing from April onwards and the final instar larvae develop during May (Rytteri et al. 2021). Since parental post-diapause larvae were exposed to temperature treatments in their final or 7^th^ instar (see next section), temperatures from May to August were used to quantify the degree of predictability separately for all 48 years.

In a time series, temporal autocorrelations measure correlations between the lagged version of itself, which allows identifying until how many days/months the variable (e.g. temperature) is predictable from the starting point. Thus, significant correlations (positive or negative) until the farthest lags will indicate that the environment is highly autocorrelated. Temporal autocorrelations were measured by creating 122 lags (123 total days from May to August) using ‘*tk_acf_diagnostics’* function from the *timetk* ver 2.6.1 package (Dancho and Vaughan 2021). It is recommended that the time series should exhibit stationarity while calculating autocorrelations and the raw temperature data did not exhibit stationarity. Thus, residuals obtained by regressing temperature values with the day number separately for each year (e.g. Shama 2015) was instead used for calculating temporal autocorrelations. Residuals had improved normality (measured using the Shapiro-Wilk test) than the raw values (Figure S1).

Finally, the SAX algorithm was used to determine the predictability of temperature. SAX is an extension of the Piecewise Aggregation Approximation (PAA) algorithm (Lin et al. 2007). PAA algorithm reduces the dimensionality of z-normalised time series while preserving important information and patterns (Keogh et al. 2001). For example, a time series of length *n* is reduced to any arbitrary length *M* where *M ≤ n* but is usually *M << n* (Keogh et al. 2001, Lin et al. 2007; see Figure 4A & B). This reduction (*n* to *M*) is carried out by dividing the original time series into *M* frames and then the mean is calculated for each frame. SAX then discretises PAA data into strings, usually alphabets (Lin et al. 2007, see Figure 4A-C). Thus, by converting continuous time series into strings of alphabets, one could investigate the similarity or repeatability of values in time series across years. Firstly, the original time series (*n=*120, see below) was reduced to 24 frames (*M*), thus one frame equals 5 days, using the ‘*paa*’ function from the *jmotif* ver 1.1.1 package (Senin 2020). PAA data was then discretised into seven alphabets from *a*-*g* (i.e. SAX conversion) using ‘*series_to_chars’* function from *jmotif* package. It is recommended to have an even number of observations in a time series while employing the SAX algorithm (see Lin et al. 2007). Thus, the 31^st^ day of May, July and August were removed to get an even number of days resulting in *n=*120 instead of 123 days for each year.

### The experiment

Diapausing larvae that were sampled in the autumn of 2020 from the Åland islands were used for the experiment. Larvae were transported to the laboratory and were kept in diapause chambers individually in Eppendorf tubes at 5°C until the start of the experiment in March 2021. Larvae were chosen from five different communes (Finstrom, Hammarland, Jomala, Saltvik and Sund) having the highest larval abundance (>200) to increase the genetic diversity of individuals in the experiment. When possible, larvae were chosen in such a way as to maximise patch-level diversity. Larval development was recommenced (n=948) by keeping them at room temperature (∼25°C) and spraying them with water. From this point, larvae were monitored for 48 hours and those who failed to show any signs of movement were considered dead. The remaining surviving larvae (n=601) were reared individually in small plastic cups and were fed daily *ad libitum* on fresh leaves of *Plantago lanceolata*.

On the day of the 5^th^ moult (i.e. mainly 6^th^ instar), the larvae were individually weighed and randomly assigned to three temperature treatments; 28°, 31° and 34°C during the day, and 9°C at night with 12L:12D cycle; in climate-controlled growth chambers (Sanyo MLR-350 and a Sanyo MLR-351). Common hostplants of *M. cinxia* (*P. lanceolata* and *Veronia spicata*) grow close to the ground and studies show that the ground temperature can be ∼10 to 20°C higher than ambient air temperature (Bennett et al. 2015, Singer & Parmesan 2018). Thus, temperatures used in our experiment are within the range of temperatures larvae experience in the field (see Verspagen et al. 2020). In the 6^th^ instar, ∼20% of larval mortality occurred due to parasitoid (*Cotesia* and *Hyposoter*) and some unknown pathogen infestation. Pupae were weighed within 24 hours and upon eclosion, adults were fed on 5% honey water and kept in their respective temperatures before they were assigned for mating. Some mortality also occurred due to failed eclosions. Males from only 28°C treatment were allowed to pair with females from all three temperatures to control for any temperature-specific paternal effects. The experimental design allows estimating both maternal and paternal effects simultaneously but carrying out an experiment of such a scale was logistically not feasible. Moreover, matings were designed to avoid pair formation between individuals from the same patches (except for one) to reduce the chances of mating between siblings. Mated females were provided with potted *P. lanceolata* plants for laying eggs and were kept in the greenhouse at 28°C.

After laying the first clutch, the total number of eggs were counted, and the clutch was split across three temperature treatments (28°, 31° & 34°C), which resulted in a full factorial split-brood design where each maternal treatment is divided into three offspring treatments (see Figure 1 for the experimental design). For three females where the number of eggs in the first clutch were low, the second clutch was also split into the same three temperature treatments. Hatched larvae were reared in groups of 15 (larvae of *M. cinxia* are gregarious in the wild) in petri-plates lined with the filter paper until diapause. Moreover, a replicate at each plate level was included to account for any plate-specific effect (Figure 1). During the second moult, three larvae were sampled from each plate for potential genomic analyses in the future. All larvae were fed daily with fresh *P. lanceolata* leaves and sprayed with water to maintain adequate humidity. Some larvae that had extremely long development times and did not show any signs of diapause were culled (n=16).

**Figure 1:**
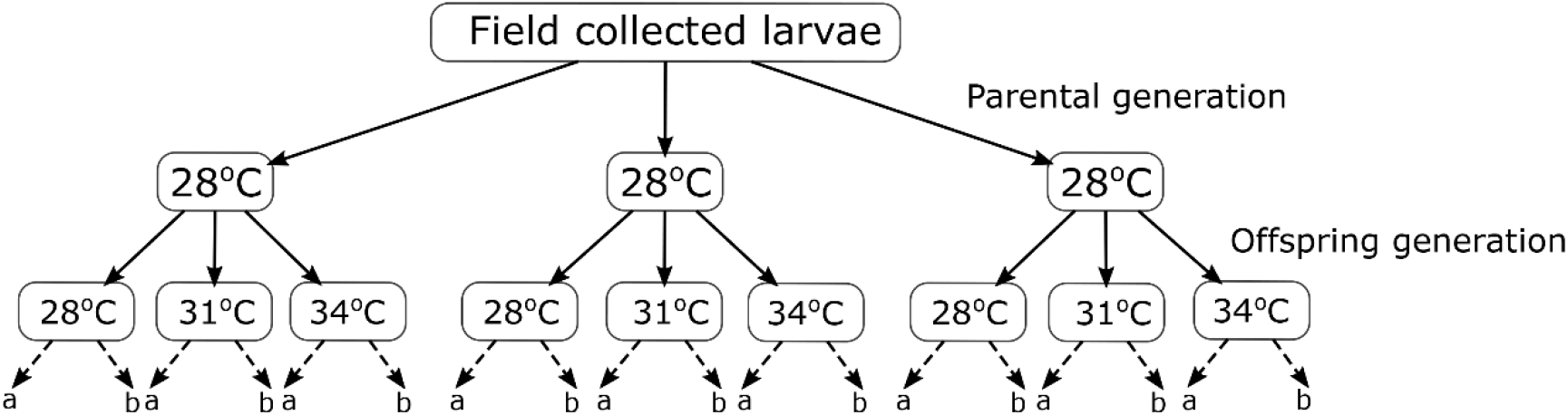
Experimental design used for estimating within- and transgenerational plasticity in the Glanville Fritillary butterfly (*Melitaea cinxia*). The dashed arrows and letters (*a & b*) at the bottom of the figure indicate replicate at the plate level (see Methods section).

### Measured life-history traits

#### Parental generation

Post-diapause larval development time (time taken from the day when diapause was broken until pupation); pupal weight; growth rate (calculated as: [ln(pupal weight)-ln(6^th^ instar weight)]/larval development time), pupal development time, life-time fecundity.

#### Offspring generation

egg development time (time taken for eggs to hatch) and hatching success; pre-diapause larval growth rate (calculated as: ln(larval weight at diapause)/pre-diapause larval development time; where larval development time is the number of days from hatching until diapause); larval survival (from 1^st^ to 3^rd^ instar & 3^rd^ instar until diapause). Larval survival was measured at two time points as three larvae were sampled at 2^nd^ moult for potential future genomic studies (see above). This approach also allows testing if survival probability differs during early and later stages of larval development.

### Statistical analyses

The effect of parental and offspring rearing temperature and their interaction on life-history traits were tested by fitting linear models (LM), linear mixed effects models (LMM) in the REML framework (*lme4* ver 1.1.27.1, Bates et al. 2015) and generalised linear mixed effects models (GLMM) (*glmmTMB* ver 1.1.2.3, Brooks et al. 2017).

#### Parental generation

LMs were fitted to test the effect of temperature, sex and their interaction on post-diapause larval development time, pupal weight, growth rate, and pupal development time (all development times were natural log-transformed). Commune (i.e. location of the larval origin) was not included as a fixed effect as removal of this term considerably improved the model fit (assessed based on the AIC score) and commune level differences in the traits are not of interest in this study. The effect of temperature on fecundity was modelled by fitting a GLMM (family = negative binomial) with clutch order (centred), temperature and their interaction as fixed effects. Pupal weight (standardized to have a mean of zero and unit standard deviation) was included as a covariate and female id or family as a random effect. Four females that laid just one clutch were removed from the analysis and egg counts up to 11 clutches were included as only a single female laid >12 clutches (16 clutches in total).

#### Offspring generation

For egg development time, LMM was fitted with parental- and offspring temperature and their interaction as fixed effects, and female id (or family) as a random effect. Similarly, egg hatching success was modelled using GLMM but specified binomial error structure with a logit link. For larval growth rate and larval survival, LMM and GLMM, respectively, were fitted with plate replicate as an additional fixed effect. Number of families across temperatures are as follows:16 or 17 at 28°C, 14 at 31°C, and 20 at 34°C.

The significance of the fixed effects and their interactions was determined based on the estimate value and their 95% confidence intervals: the effect was deemed significant if the confidence interval did not include zero. Moreover, the significance was also determined using ‘*Anova’* function from the *car* ver 3.0.11 package (Fox and Weisberg 2019) and post-hoc pairwise contrast between factors using *emmeans* ver 1.7.0 package (Lenth 2021).

## RESULTS

### The predictability of temperature fluctuations across and within seasons

When compared to the perfectly predictable photoperiod, wavelet spectra for temperature indicated that annual temperature fluctuation is overall predictable (Figure 2). That is, the spectrogram and average power plot for temperature show that there is a dominant frequency recurring at ∼365 days interval (i.e. rise and fall of temperature during summers and winters, respectively) indicated by warm colours (Figure 2). The spectrogram also shows that sharp fluctuations in temperature (indicated by white ridges) are common throughout the time series (Figure 2). Measuring temporal autocorrelations only during the growth period (from May to August) suggests that mean correlation drops rapidly with the first 10 lags and the mean value thereafter remains around zero without exceeding the confidence limits (Figure 3). However, there was high heterogeneity across years with some years showing significant negative correlations between around 30 to 70 lags (Figure 3). Thus, the temperature experienced during May is predictive of temperature during June and early July only during some years (negative correlation indicates higher temperature during June and July compared to May). Finally, comparing patterns of temperature fluctuations across years using the SAX algorithm complements the raw data that temperature generally rises during June and July (Figure 3A, Figure 4). However, the heatmap indicates that the timing of temperature rise is not synchronous across years (Figure 4D). Plotting raw values for a few years showing the highest and lowest mean temperatures during May further suggests that warmer temperatures in spring do not translate into warmer summers and *vice versa* (Figure 4E).

**Figure 2:**
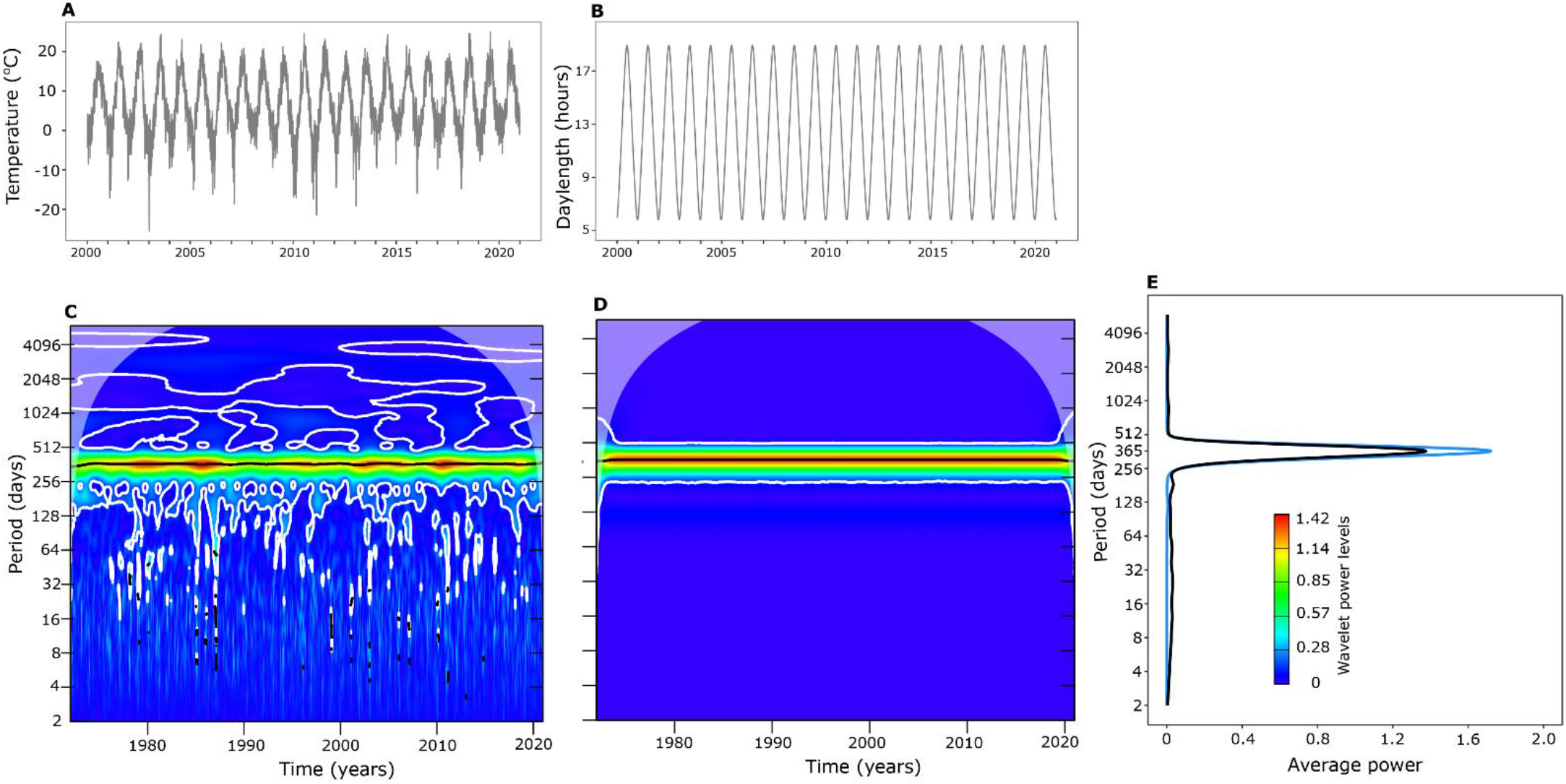
Raw time series of daily mean temperature (A) and photoperiod (B) for Jomalaby location in Åland islands and their wavelet spectra (C & D), respectively, depicting periodicity across the time series. Average power plots (E) for temperature (black curve) and photoperiod (blue curve) indicate the strength of periodicity over a particular period time period (∼ 365 days interval) for both variables. Note that in Figure A and B time series data from only 2000 to 2020 is shown for clear visualization, but wavelet analysis was performed on the entire data from 1972 to 2020. In wavelet power spectra plots, the strength of periodicity is indicated by the extent to which warmer colours are distributed over a particular time period. The black solid lines and white contour lines in spectral plots show regions of power significant at the 5% level based on 100 simulations.

**Figure 3:**
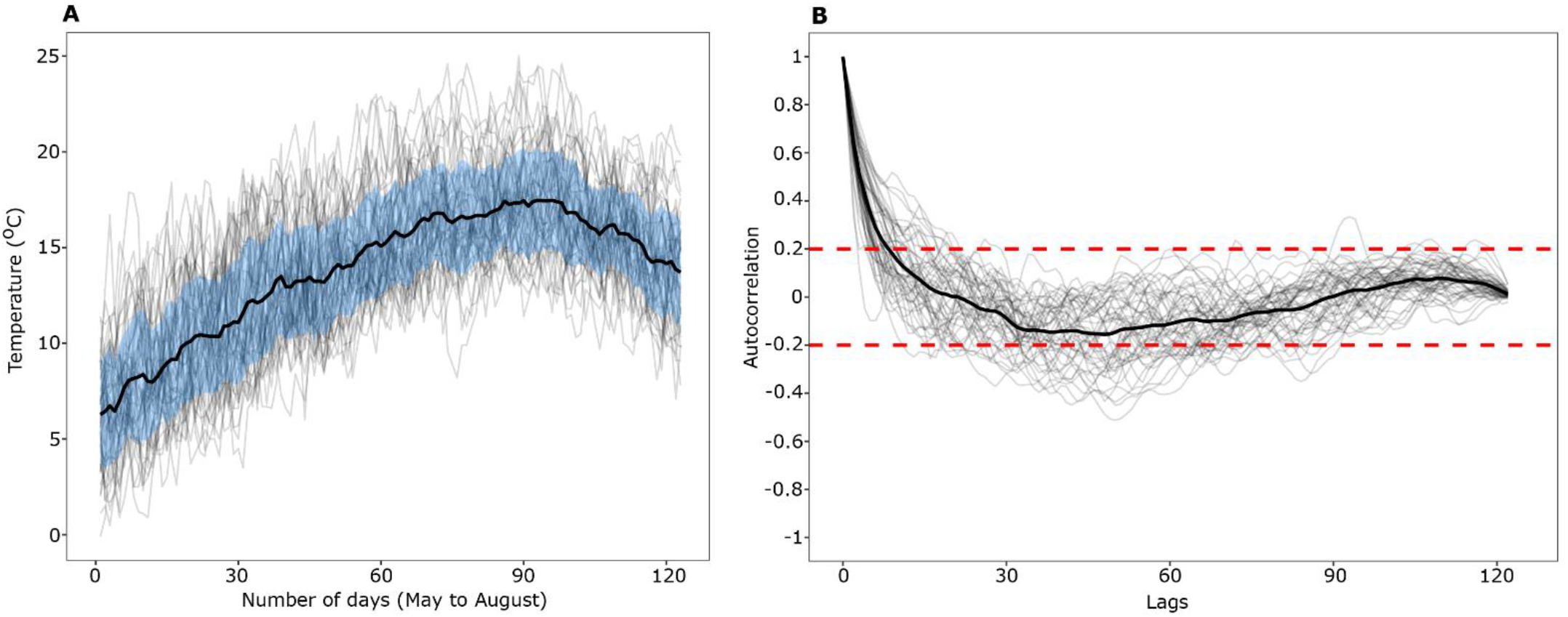
Mean daily temperatures from May-August from 1972 to 2020 (A) and their autocorrelations (B). For both plots, a thin individual line represents a single year and thick black lines indicate the mean temperature (±SD indicated by the blue band) and mean correlation (red lines indicating confidence intervals).

**Figure 4:**
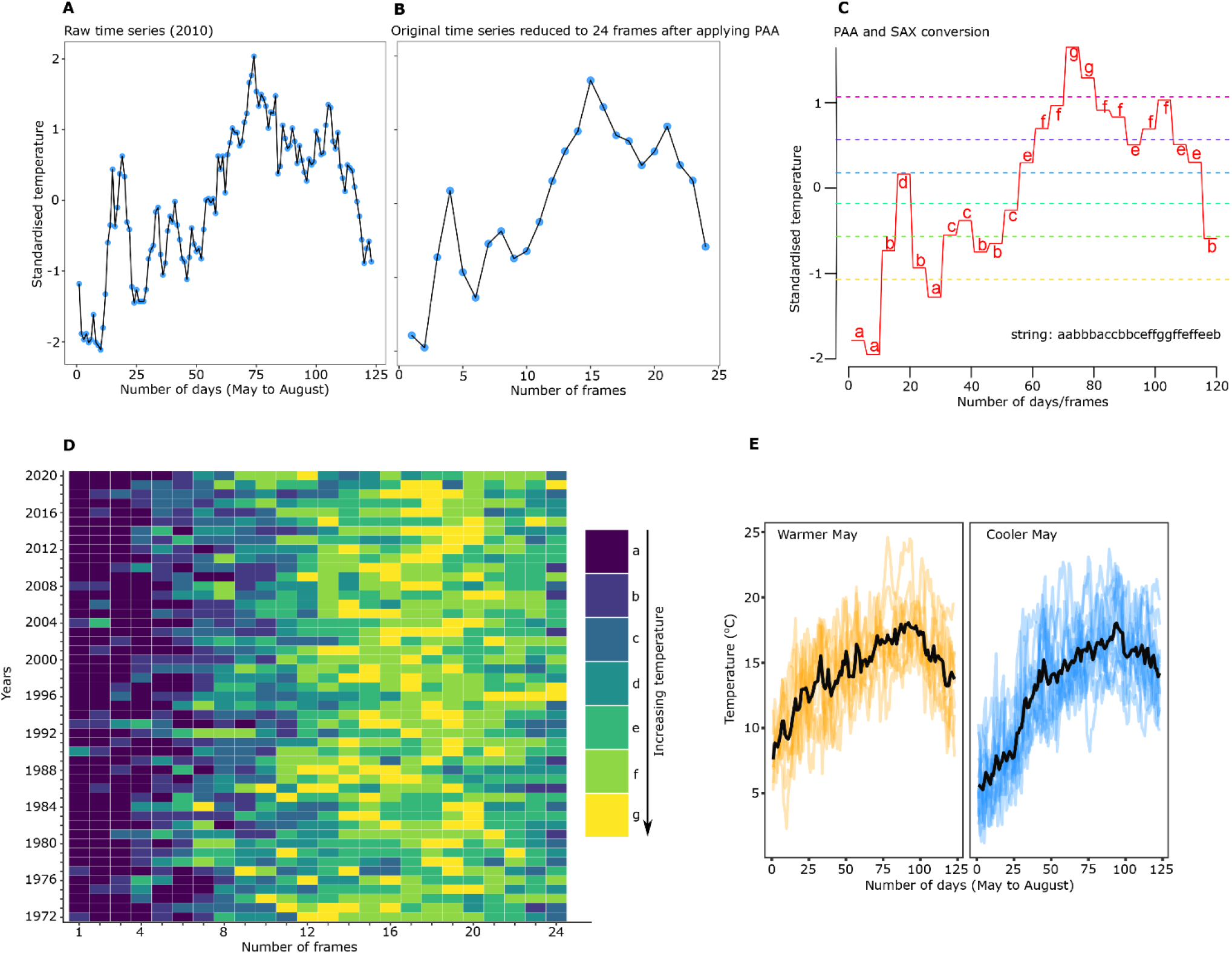
Demonstration of implementation of piecewise aggregation approximation (PAA) and symbolic aggregation approximation (SAX) algorithm on a single year (2010) from May to August (A-C). The raw time series (A, *n=*123) is reduced to the length (*M*) of 24 frames after applying the PAA algorithm (B). Figure C shows the simultaneous application of PAA and conversion of PAA data into 24 character string (i.e. SAX conversion). Obtaining such character strings across several years can then be used to investigate the repeatability or predictability of environmental variable(s) across years. The heatmap (D) was obtained after implementing the SAX algorithm for mean daily temperatures from May-August from 1972 to 2020 (here, one frame equals five days). Figure (E) shows that, for a few years which have relatively warmer (orange) and cooler (blue) temperatures in May (mean temperature indicated by thick black lines), does not necessarily translate into warmer or cooler temperatures in June or July.

### Within-generation plasticity: effect of temperature on parental life-history traits

Except for a few traits, rearing temperature during development had a strong effect on life-history traits (Table S1). Larval development times decreased while pupal weight increased with increasing temperature (Figure 5). Females developed slower and became heavier than males (Figure 5). These translated into the pattern of increasing growth rates with temperature with females having lower growth rates than males (Figure 5). Pupal development time decreased with an increase in temperature without any influence of sex (Figure S2, Table S1). The number of eggs in each clutches decreased substantially with an increasing number of clutches without any influence of developmental temperature (Figure 6). Similarly, female lifetime fecundity (calculated as the total number of eggs/number of clutches) did not differ across three developmental temperatures (One-way ANOVA, F=1.18, df=2, p=0.313, Figure). However, there was a trend that females who developed in 28°C laid fewer clutches compared to females who developed in 31° and 34°C (Figure 6).

**Figure 5:**
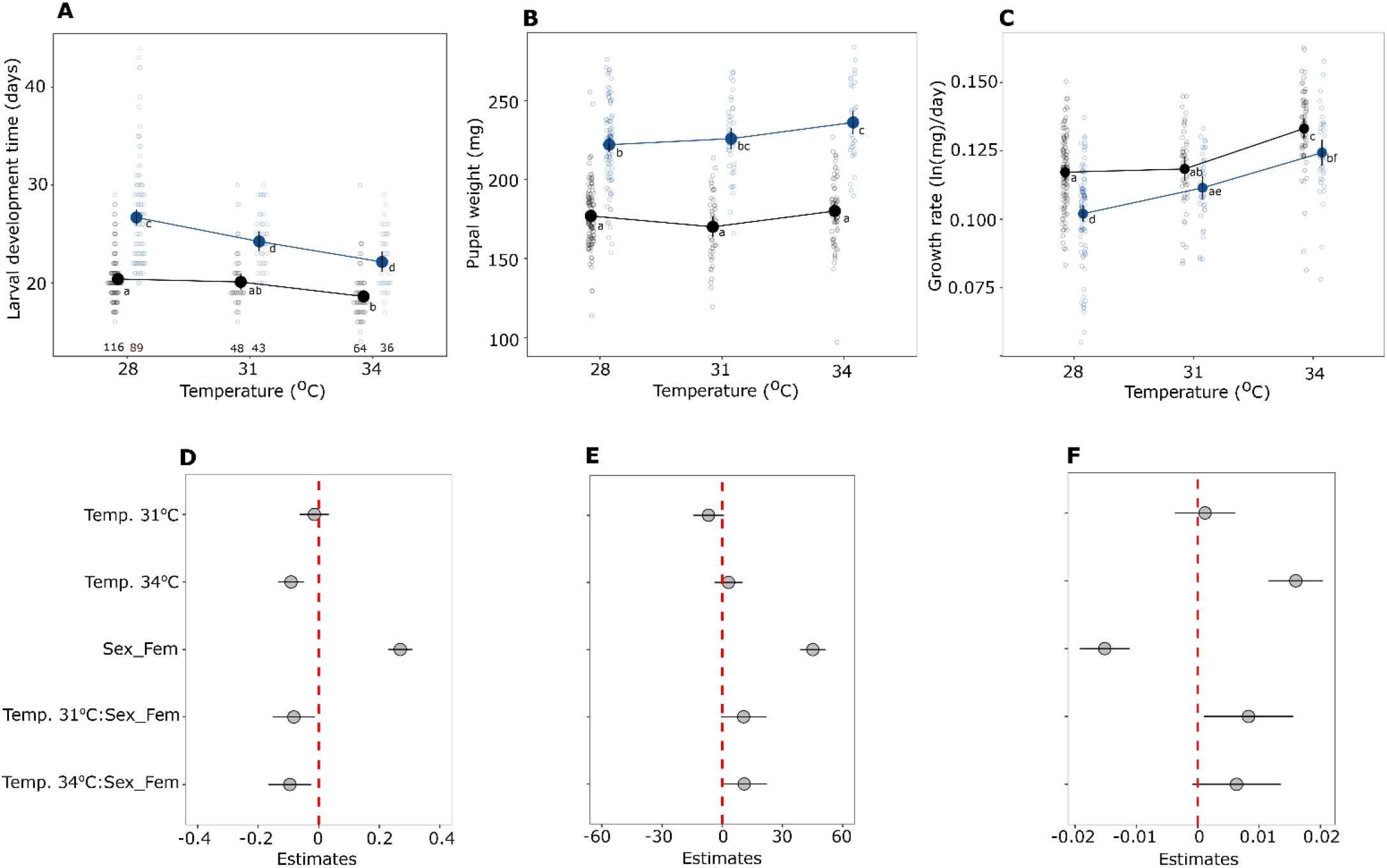
Thermal reaction norms with a model predicted mean (± 95% CI) for larval development time (A), pupal weight (B), growth rate (C), and their estimates (± 95% CI) for predictors (D-F), respectively. In Figures D-F, abbreviations are as follows: Temp. = Temperature and Sex_Fem = Sex Female. In the upper panel, sexes are denoted with different colours (blue = females, black = males) and open circles in the background represents the raw data. Significant differences between groups are indicated by different letters. Sample sizes for males and females, respectively, are provided at the bottom in Figure A and these numbers are the same for Figures B and C.

**Figure 6:**
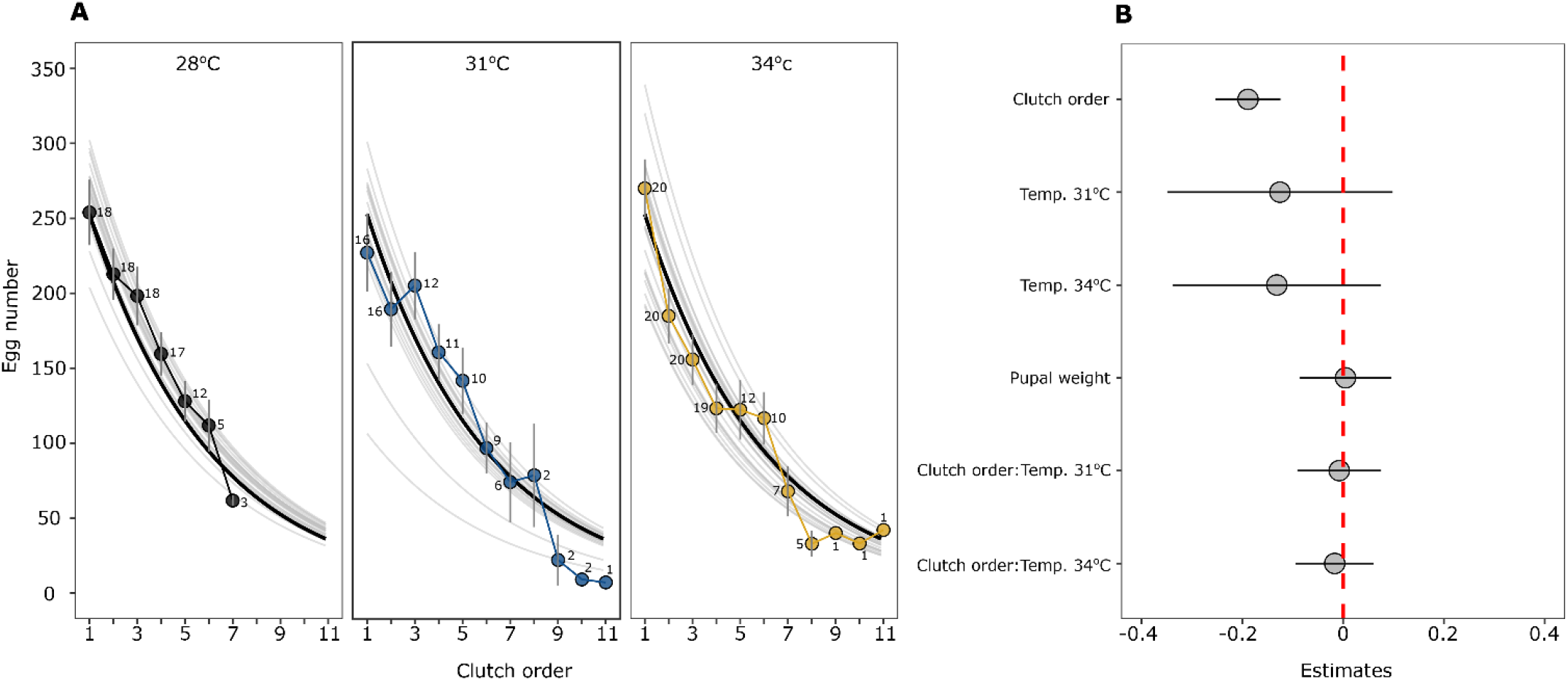
Fecundity curves across temperatures (A) and estimates (± 95% CI) for predictors (B). In Figure A, thick and thin black lines show model predicted average egg number and fecundity curves for each female, respectively, and filled circles denote average egg numbers (±SE) from the raw data. Numbers beside each circle indicate the number of females that were laying eggs.

### Transgenerational plasticity: effect of parental and offspring rearing temperature on offspring life-history traits

Under the scenario of adaptive TGP, we expected that treatments with matching parent-offspring temperatures (for example, offspring’s growing at 28°C from parents reared at 28°C) would have higher growth rates than treatments with unmatching temperatures. Egg development time decreased with increasing offspring rearing temperature without any interaction with parental temperature (Parent:Offspring, χ^2^= 6.57, df=4, p=0.15, Figure 7A & 7C, Table S2). Similarly, egg hatching success seemed to decrease with increasing offspring temperature without any interaction with parental temperature (Parent:Offspring, χ^2^= 5.79, df=4, p=0.21, Figure 7B & 7D). The post-hoc analysis further indicated that (offspring) temperature-dependent egg hatching success was statistically unclear (Figure 7A & 7B). Larval growth rates increased with increasing offspring temperature but there was also a significant interaction between parent and offspring temperature (Parent:Offspring, χ^2^= 33.32, df=4, p <0.001; Figure 8, Table S2). However, the trend was in the opposite direction than expected. That is, at 31° and 34°C offspring rearing temperature, larvae whose parents were reared at 28° had slightly higher growth rate than those from 31° and 34°, but pairwise comparisons were statistically unclear (Figure 8). Finally, larval survival from 1^st^ to 3^rd^ instar was comparatively lower at 34°C compared to 28° and 31°C offspring rearing temperature (Offspring, χ^2^= 6.78, df=2, p= 0.03; Figure 9), but there was also a weak interaction with parental temperature (Parent:Offspring, χ^2^= 11.09, df=4, p= 0.02; Figure 9). More specifically, larvae growing at 34°C from parents reared at 34°C had slightly higher survival probability than parents reared at 28°C, but not as high as those larvae from parents reared at 31°C (Figure 9). However, this effect was only observed at 34°C offspring temperature with estimates having relatively wide 95% CI and pairwise comparisons further indicated that this difference was statistically unclear (Figure 9). Such an interaction between parental and offspring temperature was absent when assessing larval survival from 3^rd^ instar until diapause (Parent:Offspring, χ^2^= 7.09, df=4, p= 0.13, Figure 9, Table S2).

**Figure 7:**
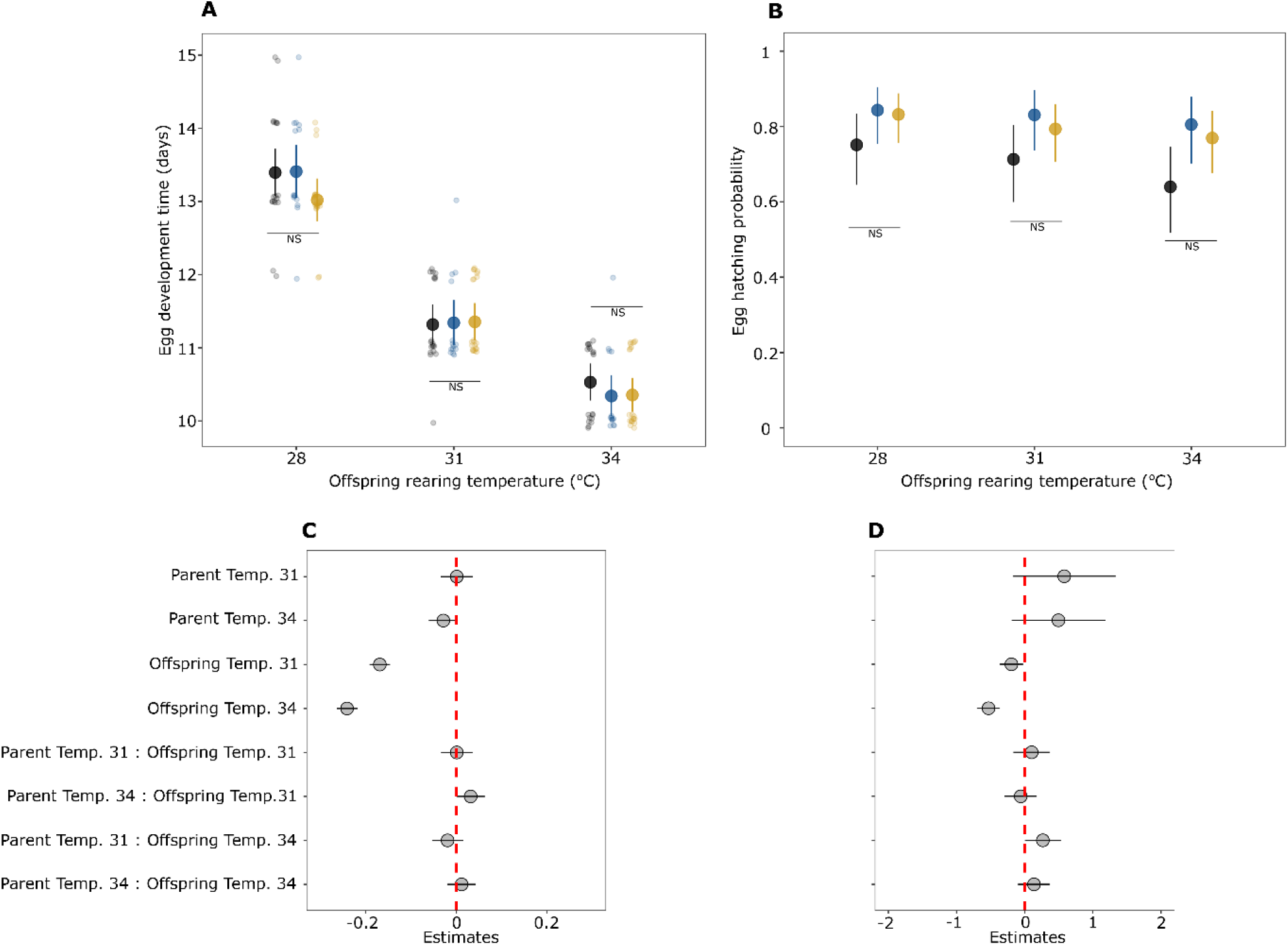
Egg development time with a model predicted mean (± 95% CI) (A) and egg hatching probability (± 95% CI) (B), and estimates of predictors (± 95% CI) for both traits, respectively, across offspring rearing temperatures. In the upper panel, coloured filled circles denote parental rearing temperatures (black = 28°C, blue = 31°C, yellow = 34°C). In Figure A, scattered points in the background show values from the raw data and NS indicates no statistical significance between pairwise contrast across parental rearing temperatures within a single offspring rearing temperature.

**Figure 8:**
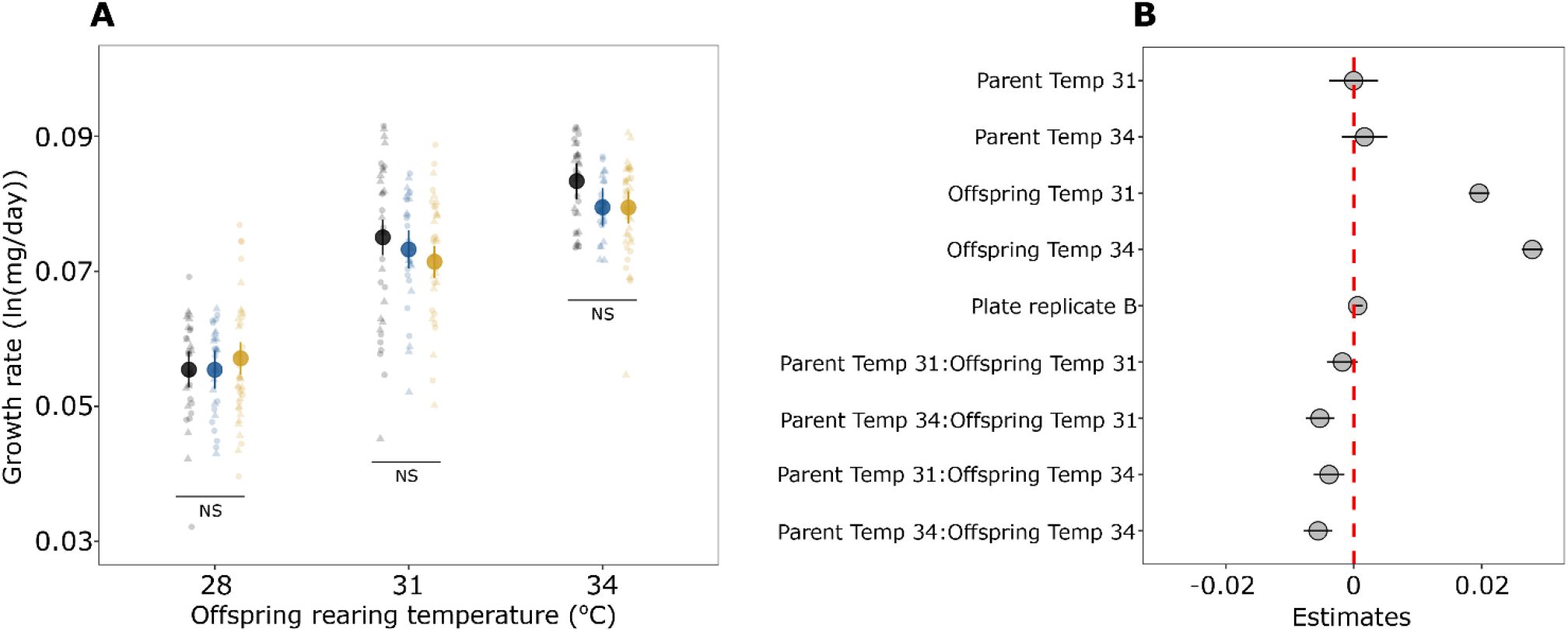
Offspring growth rates with a model predicted mean (± 95% CI) across temperatures (A) and estimates of predictors (± 95% CI) (B). In Figure A, coloured circles denote parental rearing temperatures (black = 28°C, blue = 31°C, yellow = 34°C), scattered points in the background show values from the raw data with different shapes indicating plate replicates and NS indicates no statistical significance between pairwise contrast across parental rearing temperatures within a single offspring rearing temperature.

**Figure 9:**
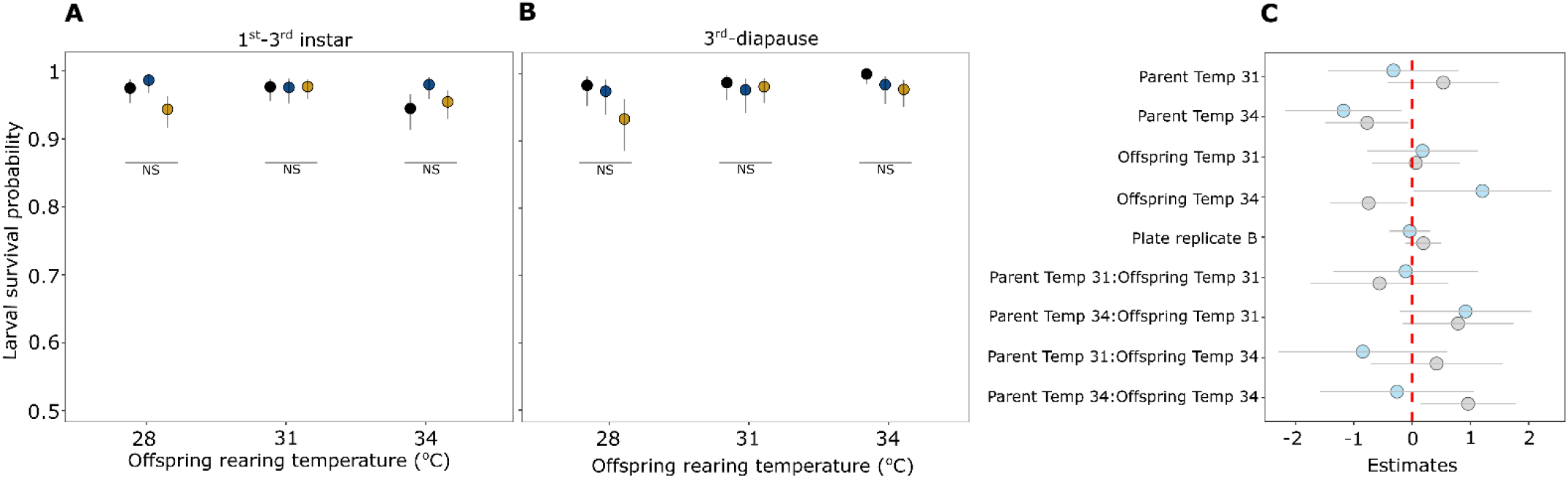
Model predicted larval survival probability (± 95% CI) from 1^st^-3^rd^ instar (A) and 3^rd^ instar until diapause (B) across offspring rearing temperatures. Figure C shows estimates (± 95% CI) for predictors with coloured circles representing survival at different time points of larval development (grey-1^st^ to 3^rd^ instar, blue – 3^rd^ instar until diapause). Coloured circles in Figure A and B denote parental rearing temperature (black = 28°C, blue = 31°C, yellow = 34°C) and NS indicates no statistical significance between pairwise contrast across parental rearing temperatures within a single offspring rearing temperature.

## DISCUSSION

The importance of TGP in adaptive evolution was recognised much earlier (e.g. Kirkpatrick and Lande 1989, Mousseau and Fox 1998, Agrawal et al. 1999) but recently there has been a renewed interest in this phenomenon even considering us to ‘rethink’ heredity (Bonduriansky 2012, Bonduriansky 2021). Moreover, even as one of the mechanisms which may allow organisms to cope with climate change (Donelson et al. 2018). Despite its widespread occurrence, renewed appreciation in adaptive evolution, and recent advancements that have unravelled the proximate basis of transgenerational inheritance (e.g. Rechavi et al. 2014), studies linking species ecology and environmental predictability to the evolution of adaptive TGP are rare. Here, using *M. cinxia* as a model system, we show that the larvae exhibit strong within-generation plasticity whereas there is only weak evidence of TGP. Furthermore, the evidence of TGP is found for two life-history traits were each in an adaptive and non-adaptive direction. Our time-series analyses further showed that, although across-season temperature fluctuations were fairly predictable, within-season fluctuations were weakly unpredictable and showed high heterogeneity in predictability (Figure 3). Based on the evidence from both temperature fluctuations and common garden experiment, we posit that our findings, at least to some extent, align with the theoretical prediction that selection will disfavour evolution of strong adaptive TGP when the predictability of offspring environment is low. We discuss our results in the context of environmental predictability and its role in the evolution of phenotypic plasticity in this butterfly, but it can also be extended to other short-lived insects in temperate regions.

When parents can predict the offspring conditions based on the prevailing environmental information, adaptive TGP can enhance offspring fitness (Burgess and Marshall 2014, Bonduriansky 2021). However, environmental predictability is scarcely quantified despite being a core pre-requisite for the evolution of adaptive TGP, especially in non-model organisms (but see Burgess and Marshall 2011, Shama 2015, Halali et al. 2021, Diaz et al. 2021). Experimental evolution studies on species with short generation times have explicitly shown that adaptive TGP readily evolves when selection lines are maintained in a temporally autocorrelated environment (e.g. Dey et al. 2016, Rescan et al. 2020, Lind et al. 2020). Few empirical studies that have quantified the predictability between parent-offspring environments have measured autocorrelations on a shorter time series (usually a few months within a year), which precludes accounting for climatic heterogeneity across years. Moreover, temporal autocorrelations suffer from a drawback that sample size decreases substantially with increasing lags and correlations for farthest lags can be unreliable. Other much informative and powerful methods such as Fourier- and wavelet transform allow quantifying environmental predictability (e.g. Burgess and Marshall 2014, Marshall and Burgess 2015, Tonkin et al. 2017, Halali et al. 2021) but are seldom used. Here we use wavelet analyses, temporal autocorrelation and the SAX algorithm to quantify the predictability of temperature fluctuations across and within the growing season as the degree of predictability at different scales can have different evolutionary implications for the evolution of phenotypic plasticity.

Given that temperature fluctuation across seasons, that is rise and fall of temperature during spring/summers and winters across 48 years is overall predictable, we propose that this pattern may have selected for the evolution of reverse temperature-size rule in this butterfly. The reverse temperature-size rule is the opposite of the more widespread pattern observed in ectotherms: the temperature-size rule (TSR) where body size decreases with increasing temperature (Atkinson 1994). Studies have demonstrated that selection can readily mould the slope of the thermal reaction norm depending on the local variation in temperatures (see Mousseau 1997). For example, a study on two populations of *Pieris rapae* butterfly demonstrated that thermal reaction norm for body size for each population exhibited negative (=TSR) and positive (= reverse-TSR) slopes (Kingsolver et al. 2007). This population-specific pattern was attributed to differences in local variation in the temperature. Populations experiencing lower and higher mean temperatures during the growing season followed the TSR and reverse-TSR, respectively (Kingsolver et al. 2007). Such local adaptation in thermal reaction norm is likely to be adaptive as several life-history traits co-vary with body size. For example, the larger body size is correlated with increased longevity, higher survival success during harsh periods and increased fecundity (Honěk 1993, Blanckenhorn 2000).

The raw temperatures indicate a consistent trend of gradual increase of temperature from May to July although there is a lot of variation across years (Figure). Since post-diapause larvae grow from April to May and adults eclose during early June, the mean temperature experienced by later stages of the development (i.e. 7^th^ instar larvae and pupae) is higher compared to the earlier 6^th^ instar larvae. Thus, selection may have favoured the thermal reaction norm with a positive slope (i.e. reverse-TSR) such that a larger body size is achieved even when the temperatures are higher (e.g. Kingsolver et al. 2007). Interestingly, the temperature did not have a strong effect on fecundity despite females having larger body sizes (pupal weight as a proxy) at higher temperatures, but this correlation was apparent in another study where fecundity was assessed under semi-natural field conditions (see Rosa and Saastamoinen 2017). Also, there was a trend that females that developed under 31°C and 34°C laid more clutches than those developed at 28°C (Figure 6). We posit that since this increase in temperature during spring/summer is overall predictable across years (based on wavelet analysis), such a pattern may have favoured the evolution of reverse-TSR in this butterfly, and likely in other univoltine temperate insects. In contrast to post-diapause larvae, early stages of pre-diapause larvae experience higher temperatures than later stages as the temperature gradually decreases from July to August. Links between temperature and body size is not as straightforward in pre-diapause larvae as larvae can diapause either in the 4^th^ or 5^th^ instar and can show large differences in body mass at diapause (Saastamoinen et al. 2013a, Kahilainen et al. 2021).

Fine-scale variation of temperature and its predictability within the growing season can have important consequences for the evolution of TGP. Temporal autocorrelations show that average correlations are near zero suggesting temperatures during May do not or are extremely weakly predictive of temperatures in succeeding months (Figure 3). Moreover, the heatmap derived from the SAX algorithm (Figure 4D) suggests that, as expected, June and July are relatively warmer, but the timing of temperature peaks are not synchronous across years. Examining raw temperatures further shows that years having relatively warmer temperatures during May does not necessarily translate into a warmer summer (i.e. June and July, Figure 4E). This, therefore, indicates that temperatures experienced by parents during their growth are not or are only weakly predictive of the temperature that will be experienced by the offspring. Overall, we posit that evidence of weak adaptive TGP in our system may have been due to lower predictability of temperature within the growing season and selection may have instead favoured the evolution of strong within-generation plasticity which provides a more rapid response to the prevailing conditions.

Interestingly, there was an interaction between parental and offspring temperature for two offspring life-history traits (pre-diapause larval growth rate and survival), but both had low effect sizes with relatively wide confidence intervals. We found that larvae whose mothers were reared at 28°C had slightly higher growth rates at warmer, 31°C and 34°C, offspring rearing temperature than larvae from mothers reared warmer temperatures (Figure). This pattern is, however, in the opposite direction than we initially expected (i.e. that offspring’s growing in similar temperatures as that of their parents would have higher performance). We speculate that this trend may still be adaptive, as 28°C is not optimal for achieving high growth rates, and thus parents who developed at 28°C may prime their offspring’s to attain higher growth rates in warmer conditions. Furthermore, there was weak evidence for TGP towards the expected direction for offspring survival from 1^st^ to 3^rd^ instar. That is, at 34°C offspring rearing temperature, offspring whose parents were reared at higher temperature (31° & 34°C) had slightly higher survival (Figure). However, this weak effect was only observed at 34°C offspring rearing temperature and this trend was absent when larval survival was measured from the 3^rd^ instar until diapause. Moreover, there was an indication that, especially when estimating fitness in terms of offspring survival, 34°C appeared to be the stressful condition. Theoretically, it is argued that TGP is not always adaptive, in fact, in most instances, it is expected to be mal- or non-adaptive (Bonduriansky 2021). TGP may occur from factors other than adaptive reasons such as high physiological sensitivity to environmental fluctuations during reproduction or physiological constraints (Bonduriansky 2021). Significant parent-offspring interactions for two life-history traits in this study and evidence from other studies in *M. cinxia* (Saastamoinen et al. 2013b, Salgado and Saastamoinen 2019) does suggest that TGP is prevalent in this system. However, whether the TGP is actually adaptive will require new ways of measuring fitness in both the laboratory and wild.

Commonly used hostplants of *M. cinxia, Plantago lanceolata* and *Veronica spicata*, grow close to the ground and studies have shown that ground temperature can be ∼10-20°C degrees warmer than ambient air temperature (Bennett et al. 2015, Singer and Parmesan 2018). We acknowledge that using time series data on the ground temperature in microhabitats of *M. cinxia* would have been ideal, but such data is not yet available. However, studies have suggested that there is generally a strong correlation between air and ground temperature (Tsilingiridis and Papakostas 2014). Moreover, during exceptionally warm years such as in 2018 when summer temperatures were >25° in Åland (see van bergen et al. 2020), ground temperatures may have easily exceeded 34°C which was the warmest treatment in our experiment. Future studies using higher stress-inducing temperatures in the experiments will be important in investigating the prevalence of adaptive TGP.

One of the notable effects of climate change, especially at the higher latitudes, is that the springs are arriving early, and summers are getting longer (Bradshaw & Holzapfel 2001). Climate change is also predicted to increase temporal and spatial autocorrelation of temperature in temperate regions (Di Cecco and Gouhier 2018, Kahilainen et al. 2018). Studies in both plants and animals have indicated that TGP might be one of the mechanisms enabling species to buffer the effects of climate change (e.g. Groot et al. 2017). Given that we find very weak evidence for adaptive TGP, it will be of great interest to investigate if the species could evolve such a response on a contemporary time scale. Future studies using simulated climate change experiments (e.g. Shama et al. 2014) would allow testing such a hypothesis. Moreover, performing experiments similar to the current study on *M. cinxia* populations from lower latitudes (e.g. from France, Spain, Morocco; similar to Munch et al. 2021), where temporal autocorrelations for temperature are expected to be higher would allow investigating how climatic predictability drives the evolution adaptive TGP.

## AUTHOR CONTRIBUTIONS

SH & MS conceived the idea and designed the experiment; SH carried out the experiment, performed analyses, and wrote the manuscript with inputs from MS.

## DATA ACCESSIBILITY STATEMENT

All raw data will be made available in a public repository upon acceptance of the manuscript.

## ACKNOWLEDGEMENTS

We thank Nadja Verspagen, Annina Rokka, Mikko Immonen, Jefferson Delgado Florez, Morganne Ducoin, Basile Marteau, and Li Guohua, for their assistance during the experiment. SH also thanks Charles Rocabert for general discussions on time series analysis and for recommending the SAX algorithm. This work was supported by a start-up grant from HiLIFE to MS.

## REFERENCES

1. Agrawal, A. A., Laforsch, C., & Tollrian, R. (1999). Transgenerational induction of defences in animals and plants. Nature, 401(6748), 60–63.

2. Atkinson, D. (1994). Temperature and organism size: a biological law for ectotherms?. Advances in Ecological Research, 25, 1–58.

3. Bates, D., Maechler, M., Bolker, B., & Walker, S. (2015). Fitting Linear Mixed-Effects Models Using lme4. Journal of Statistical Software, 67(1), 1–48.

4. Beldade, P., Mateus, A. R. A., & Keller, R. A. (2011). Evolution and molecular mechanisms of adaptive developmental plasticity. Molecular Ecology, 20(7), 1347–1363.

5. Bell, A. M., & Hellmann, J. K. (2019). An integrative framework for understanding the mechanisms and multigenerational consequences of transgenerational plasticity. Annual Review of Ecology, Evolution, and Systematics, 50, 97–118.

6. Bennett, N. L., Severns, P. M., Parmesan, C., & Singer, M. C. (2015). Geographic mosaics of phenology, host preference, adult size and microhabitat choice predict butterfly resilience to climate warming. Oikos, 124(1), 41–53.

7. Bivand, R., & Lewin-Koh, N. (2021). maptools: Tools for Handling Spatial Objects. R package version 1.1-2.

8. Blanckenhorn, W. U. (2000). The evolution of body size: what keeps organisms small?. The Quarterly Review of Biology, 75(4), 385–407.

9. Bonamour, S., Chevin, L. M., Charmantier, A., & Teplitsky, C. (2019). Phenotypic plasticity in response to climate change: the importance of cue variation. Philosophical Transactions of the Royal Society B, 374(1768), 20180178.

10. Bonduriansky, R. (2012). Rethinking heredity, again. Trends in Ecology & Evolution, 27(6), 330–336.

11. Bonduriansky, R. (2021). Plasticity across generations. In Phenotypic Plasticity & Evolution (pp. 327–348). CRC Press.

12. Bradshaw, W. E., & Holzapfel, C. M. (2001). Genetic shift in photoperiodic response correlated with global warming. Proceedings of the National Academy of Sciences, 98(25), 14509–14511.

13. Brooks, M. E., Kristensen, K., Van Benthem, K. J., Magnusson, A., Berg, C. W., Nielsen, A., Skaug, H.J., Martin, M., & Bolker, B. M. (2017). glmmTMB balances speed and flexibility among packages for zero-inflated generalized linear mixed modeling. The R journal, 9(2), 378–400.

14. Burgess, S. C., & Marshall, D. J. (2011). Temperature-induced maternal effects and environmental predictability. Journal of Experimental Biology, 214(14), 2329–2336.

15. Burgess, S. C., & Marshall, D. J. (2014). Adaptive parental effects: the importance of estimating environmental predictability and offspring fitness appropriately. Oikos, 123(7), 769–776.

16. Cazelles, B., Cazelles, K., & Chavez, M. (2014). Wavelet analysis in ecology and epidemiology: impact of statistical tests. Journal of the Royal Society Interface, 11(91), 20130585.

17. Cazelles, B., Chavez, M., Berteaux, D., Ménard, F., Vik, J. O., Jenouvrier, S., & Stenseth, N. C. (2008). Wavelet analysis of ecological time series. Oecologia, 156(2), 287–304.

18. Chevin, L. M., & Lande, R. (2015). Evolution of environmental cues for phenotypic plasticity. Evolution, 69(10), 2767–2775.

19. Colicchio, J. M., & Herman, J. (2020). Empirical patterns of environmental variation favor adaptive transgenerational plasticity. Ecology and Evolution, 10(3), 1648–1665.

20. Dancho, M., & Vaughan, D. (2021). timetk: A Tool Kit for Working with Time Series in R. R package version 2.6.1.

21. Dey, S., Proulx, S. R., & Teotonio, H. (2016). Adaptation to temporally fluctuating environments by the evolution of maternal effects. PLoS Biology, 14(2), e1002388.

22. Di Cecco, G. J., & Gouhier, T. C. (2018). Increased spatial and temporal autocorrelation of temperature under climate change. Scientific Reports, 8(1), 1–9.

23. Diaz, F., Kuijper, B., Hoyle, R. B., Talamantes, N., Coleman, J. M., & Matzkin, L. M. (2021). Environmental predictability drives adaptive within‐and transgenerational plasticity of heat tolerance across life stages and climatic regions. Functional Ecology, 35(1), 153–166.

24. Donelson, J. M., Salinas, S., Munday, P. L., & Shama, L. N. (2018). Transgenerational plasticity and climate change experiments: where do we go from here?. Global Change Biology, 24(1), 13–34.

25. Fox, C. W., Thakar, M. S., & Mousseau, T. A. (1997). Egg size plasticity in a seed beetle: an adaptive maternal effect. The American Naturalist, 149(1), 149–163.

26. Fox, J., & Weisberg, S. (2019). An R Companion to Applied Regression, Third edition. Sage, Thousand Oaks CA.

27. Galloway, L. F., & Etterson, J. R. (2007). Transgenerational plasticity is adaptive in the wild. Science, 318(5853), 1134–1136.

28. Groot, M. P., Kubisch, A., Ouborg, N. J., Pagel, J., Schmid, K. J., Vergeer, P., & Lampei, C. (2017). Transgenerational effects of mild heat in Arabidopsis thaliana show strong genotype specificity that is explained by climate at origin. New Phytologist, 215(3), 1221–1234.

29. Halali, S., Halali, D., Barlow, H. S., Molleman, F., Kodandaramaiah, U., Brakefield, P. M., & Brattström, O. (2021). Predictability of temporal variation in climate and the evolution of seasonal polyphenism in tropical butterfly communities. Journal of Evolutionary Biology, 34, 1362–1375.

30. Honěk, A. (1993). Intraspecific variation in body size and fecundity in insects: a general relationship. Oikos, 66(3), 483–492.

31. Jablonka, E. (2017). The evolutionary implications of epigenetic inheritance. Interface Focus, 7(5), 20160135.

32. Kahilainen, A., Oostra, V., Somervuo, P., Minard, G., & Saastamoinen, M. (2021). Alternative developmental and transcriptomic responses to host plant water limitation in a butterfly metapopulation. Molecular Ecology. In Press

33. Kahilainen, A., van Nouhuys, S., Schulz, T., & Saastamoinen, M. (2018). Metapopulation dynamics in a changing climate: Increasing spatial synchrony in weather conditions drives metapopulation synchrony of a butterfly inhabiting a fragmented landscape. Global Change Biology, 24(9), 4316–4329.

34. Keogh, E., Chakrabarti, K., Pazzani, M., & Mehrotra, S. (2001). Dimensionality reduction for fast similarity search in large time series databases. Knowledge and Information Systems, 3(3), 263–286.

35. Kingsolver, J. G., & Huey, R. B. (2008). Size, temperature, and fitness: three rules. Evolutionary Ecology Research, 10(2), 251–268.

36. Kingsolver, J. G., Massie, K. R., Ragland, G. J., & Smith, M. H. (2007). Rapid population divergence in thermal reaction norms for an invading species: breaking the temperature–size rule. Journal of Evolutionary Biology, 20(3), 892–900.

37. Kirkpatrick, M., & Lande, R. (1989). The evolution of maternal characters. Evolution, 43(3), 485–503.

38. Leimar, O., & McNamara, J. M. (2015). The evolution of transgenerational integration of information in heterogeneous environments. The American Naturalist, 185(3), E55–E69.

39. Lenth, R.V. (2021). emmeans: Estimated Marginal Means, aka Least-Squares Means. R package version 1.7.0.

40. Levis, N. A., & Pfennig, D. W. (2016). Evaluating ‘plasticity-first’ evolution in nature: key criteria and empirical approaches. Trends in Ecology & Evolution, 31(7), 563–574.

41. Lin, J., Keogh, E., Wei, L., & Lonardi, S. (2007). Experiencing SAX: a novel symbolic representation of time series. Data Mining and Knowledge Discovery, 15(2), 107–144.

42. Lind, M. I., Zwoinska, M. K., Andersson, J., Carlsson, H., Krieg, T., Larva, T., & Maklakov, A. (2020). Environmental variation mediates the evolution of anticipatory parental effects. Evolution Letters, 4(4), 371–381.

43. Ma, C. S., Ma, G., & Pincebourde, S. (2021). Survive a warming climate: insect responses to extreme high temperatures. Annual Review of Entomology, 66, 163–184.

44. Marshall, D. J., & Burgess, S. C. (2015). Deconstructing environmental predictability: seasonality, environmental colour and the biogeography of marine life histories. Ecology Letters, 18(2), 174–181.

45. Moran, N. A. (1992). The evolutionary maintenance of alternative phenotypes. The American Naturalist, 139(5), 971–989.

46. Mousseau, T. A. (1997). Ectotherms follow the converse to Bergmann’s rule. Evolution, 51(2), 630–632.

47. Mousseau, T. A., & Fox, C. W. (1998). The adaptive significance of maternal effects. Trends in Ecology & Evolution, 13(10), 403–407.

48. Munch, S. B., Lee, W. S., Walsh, M., Hurst, T., Wasserman, B. A., Mangel, M., & Salinas, S. (2021). A latitudinal gradient in thermal transgenerational plasticity and a test of theory. Proceedings of the Royal Society B, 288(1950), 20210797.

49. Ovaskainen, O., & Saastamoinen, M. (2018). Frontiers in metapopulation biology: The legacy of Ilkka Hanski. Annual Review of Ecology, Evolution, and Systematics, 49, 231–252.

50. Pfennig, D. W., Wund, M. A., Snell-Rood, E. C., Cruickshank, T., Schlichting, C. D., & Moczek, P. (2010). Phenotypic plasticity’s impacts on diversification and speciation. Trends in Ecology & Evolution, 25(8), 459–467.

51. Pigliucci, M. (2005). Evolution of phenotypic plasticity: where are we going now?. Trends in Ecology & Evolution, 20(9), 481–486.

52. R Core Team (2021). R: A language and environment for statistical computing. R Foundation for Statistical Computing, Vienna, Austria.

53. Rechavi, O., Houri-Ze’evi, L., Anava, S., Goh, W. S. S., Kerk, S. Y., Hannon, G. J., & Hobert, O. (2014). Starvation-induced transgenerational inheritance of small RNAs in C. elegans. Cell, 158(2), 277–287.

54. Reed, T. E., Waples, R. S., Schindler, D. E., Hard, J. J., & Kinnison, M. T. (2010). Phenotypic plasticity and population viability: the importance of environmental predictability. Proceedings of the Royal Society B, 277(1699), 3391–3400.

55. Rescan, M., Grulois, D., Ortega-Aboud, E., & Chevin, L. M. (2020). Phenotypic memory drives population growth and extinction risk in a noisy environment. Nature Ecology & Evolution, 4(2), 193–201.

56. Rosa, E., & Saastamoinen, M. (2017). Sex-dependent effects of larval food stress on adult performance under semi-natural conditions: only a matter of size?. Oecologia, 184(3), 633–642.

57. Rösch, A., & Schmidbauer, H. (2018a). WaveletComp 1.1: A guided tour through the R package.

58. Rösch, A., & Schmidbauer, H. (2018b). WaveletComp: Computational Wavelet Analysis. R package version 1.1.

59. Rytteri, S., Kuussaari, M., & Saastamoinen, M. (2021). Microclimatic variability buffers butterfly populations against increased mortality caused by phenological asynchrony between larvae and their host plants. Oikos, 130(5), 753–765.

60. Saastamoinen, M., Hirai, N., & van Nouhuys, S. (2013b). Direct and trans-generational responses to food deprivation during development in the Glanville fritillary butterfly. Oecologia, 171(1), 93–104.

61. Saastamoinen, M., Ikonen, S., Wong, S. C., Lehtonen, R., & Hanski, I. (2013a). Plastic larval development in a butterfly has complex environmental and genetic causes and consequences for population dynamics. Journal of Animal Ecology, 82(3), 529–539.

62. Salgado, A. L., & Saastamoinen, M. (2019). Developmental stage-dependent response and preference for host plant quality in an insect herbivore. Animal Behaviour, 150, 27–38.

63. Salinas, S., & Munch, S. B. (2012). Thermal legacies: transgenerational effects of temperature on growth in a vertebrate. Ecology Letters, 15(2), 159–163.

64. Senin, P. (2020). jmotif: Time Series Analysis Toolkit Based on Symbolic Aggregate Discretization, i.e. SAX. R package version 1.1.1.

65. Shama, L. N. (2015). Bet hedging in a warming ocean: predictability of maternal environment shapes offspring size variation in marine sticklebacks. Global Change Biology, 21(12), 4387–4400.

66. Shama, L. N., Strobel, A., Mark, F. C., & Wegner, K. M. (2014). Transgenerational plasticity in marine sticklebacks: maternal effects mediate impacts of a warming ocean. Functional Ecology, 28(6), 1482–1493.

67. Shapiro, A. M. (1976). Seasonal polyphenism. In Evolutionary Biology (}xpp. 259–333). Springer, Boston, MA.

68. Singer, M. C., & Parmesan, C. (2018). Lethal trap created by adaptive evolutionary response to an exotic resource. Nature, 557(7704), 238–241.

69. Taborsky, B., English, S., Fawcett, T. W., Kuijper, B., Leimar, O., McNamara, J. M., Ruuskanen, S., & Sandi, C. (2021). Towards an evolutionary theory of stress responses. Trends in Ecology & Evolution, 36(1), 39–48.

70. Tonkin, J. D., Bogan, M. T., Bonada, N., Rios‐Touma, B., & Lytle, D. A. (2017). Seasonality and predictability shape temporal species diversity. Ecology, 98(5), 1201–1216.

71. Tsilingiridis, G., & Papakostas, K. (2014). Investigating the relationship between air and ground temperature variations in shallow depths in northern Greece. Energy, 73, 1007–1016.

72. Uller, T. (2008). Developmental plasticity and the evolution of parental effects. Trends in Ecology & Evolution, 23(8), 432–438.

73. Uller, T., Nakagawa, S., & English, S. (2013). Weak evidence for anticipatory parental effects in plants and animals. Journal of Evolutionary Biology, 26(10), 2161–2170.

74. van Bergen, E., Dallas, T., DiLeo, M. F., Kahilainen, A., Mattila, A. L., Luoto, M., & Saastamoinen, M. (2020). The effect of summer drought on the predictability of local extinctions in a butterfly metapopulation. Conservation Biology, 34(6), 1503–1511.

75. Verspagen, N., Ikonen, S., Saastamoinen, M., & van Bergen, E. (2020). Multidimensional plasticity in the Glanville fritillary butterfly: larval performance is temperature, host and family specific. Proceedings of the Royal Society B, 287(1941), 20202577.

76. Wahlberg, N. (2000). Comparative descriptions of the immature stages and ecology of five Finnish melitaeine butterfly species (Lepidoptera: Nymphalidae). Entomologica Fennica, 11(3), 167–174.

77. Wickham, H. (2016). ggplot2: elegant graphics for data analysis. Springer.

78. Wickham, H., Averick, M., Bryan, J., Chang, W., McGowan, L. D. A., François, R., et al. (2019). Welcome to the Tidyverse. Journal of Open Source Software, 4(43), 1686.

79. Yin, J., Zhou, M., Lin, Z., Li, Q. Q., & Zhang, Y. Y. (2019). Transgenerational effects benefit offspring across diverse environments: A meta‐analysis in plants and animals. Ecology Letters, 22(11), 1976–1986.

80. Zeileis, A., & Grothendieck, G. (2005). zoo: S3 Infrastructure for Regular and Irregular Time Series. Journal of Statistical Software, 14(6), 1–27.

81. Zizzari, Z. V., & Ellers, J. (2014). Rapid shift in thermal resistance between generations through maternal heat exposure. Oikos, 123(11), 1365–1370.

