## Supplementary information file for "Exploring links between climatic predictability and the evolution of within- and transgenerational plasticity"

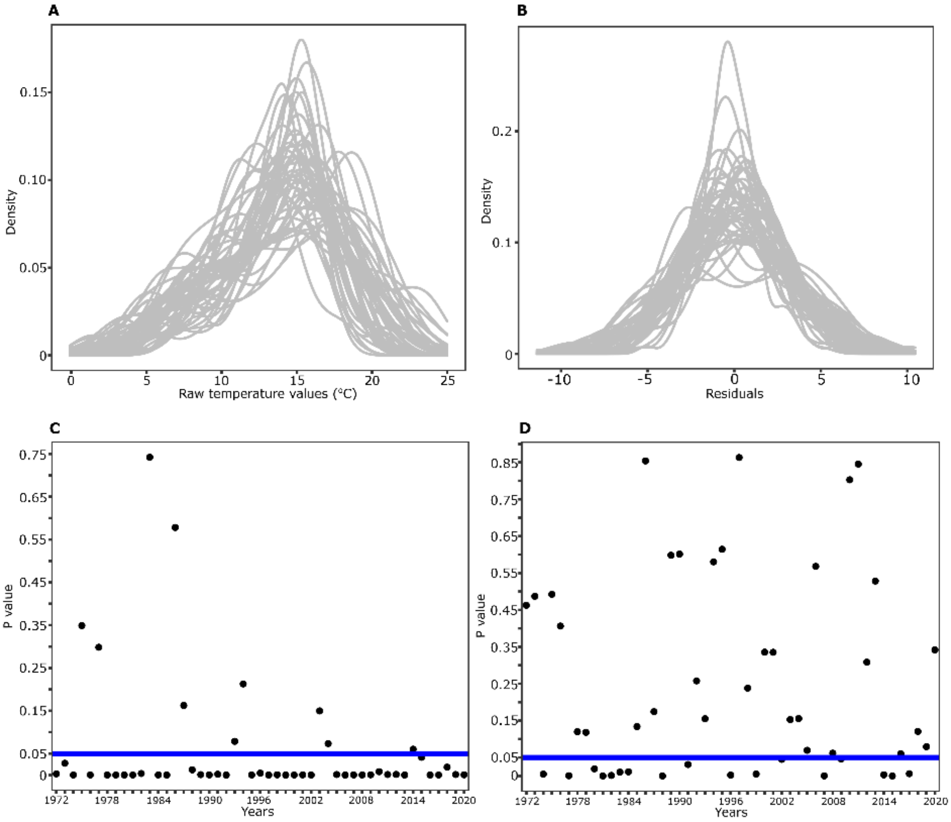


**Figure S1:** Distribution of raw temperature values and their residuals (obtained by performing regression with day number separately for each year) from May to August for the entire time series from 1972 to 2020. A single curve in Figures A and B represents a single year. Figures C and D depict P values for each year obtained from the Shapiro-Wilk test to test the normality of the time series.

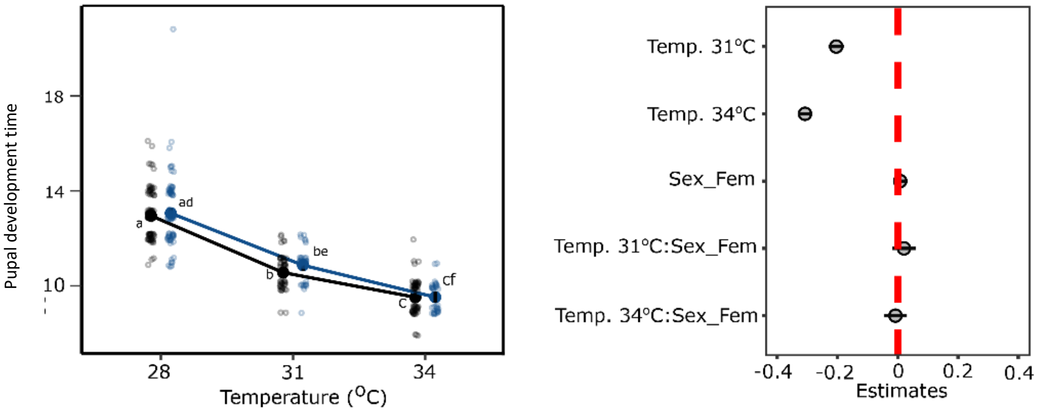


**Figure S2:** Thermal reaction norm with a mean (± 95% CI) for pupal development time (A) and estimates (± 95% CI) for predictors (B). In Figure A, sexes are denoted with different colours (blue - females, black – males) and open circles in the background represents the raw data. Significant differences between groups are indicated by different letters

**Table S1:** Anova table obtained after fitting linear models (LM) and generalized linear mixed models (GLMM) for several life history traits in the parental generation. Please refer to the figures in the main text for estimate values for predictors.

| Trait | Term | Sumsq | df | Statistic | P value |
|---|---|---|---|---|---|
| *Larval development time (LM: ~ Temperature * sex)* | | | | | |
| | Temperature | 1.121902 | 2 | 27.8198 | <0.001 |
| | Sex | 4.950894 | 1 | 245.5346 | <0.001 |
| | Temperature:Sex | 0.191293 | 2 | 4.743494 | 0.0092 |
| *Pupal weight (LM: ~ Temperature * sex)* | | | | | |
| | Temperature | 5088.6551 | 2 | 4.8609 | 0.0082 |
| | Sex | 242575.5995 | 1 | 463.4356 | <0.001 |
| | Temperature:Sex | 2795.8604 | 2 | 2.6707 | 0.0705 |
| *Growth rate (LM: ~ Temperature * sex)* | | | | | |
| | Temperature | 0.02305 | 2 | 54.0780 | <0.001 |
| | Sex | 0.01314 | 1 | 61.6344 | <0.001 |
| | Temperature:Sex | 0.00132 | 2 | 3.0943 | 0.0464 |
| *Eclosion time (LM: ~ Temperature * sex)* | | | | | |
| | Temperature | 7.0311 | 2 | 593.6177 | <0.001 |
| | Sex | 0.0122 | 1 | 2.0578 | 0.1522 |
| | Temperature:Sex | 0.0102 | 2 | 0.8598 | 0.4240 |
| *Fecundity (GLMM: ~ clutch order * Temperature + pupal weight + (1\|family))* | | | | | |
| | Clutch order | | 1 | 174.8144 | <0.001 |
| | Temperature | | 2 | 1.737972 | 0.4193 |
| | Pupal weight | | 1 | 0.011122 | 0.9160 |
| | Clutch order:Temperature | | 2 | 0.189759 | 0.9094 |

**Table S2:** Anova table obtained after fitting linear mixed models (LMM) and generalized linear mixed models (GLMM) for several life history traits in the offspring generation. Please refer to the figures in the main text for estimate values for predictors.

| Trait | Term | df | Statistic | P value |
|---|---|---|---|---|
| *Egg development time (LMM: ~ Parent temp * Offspring temp + (1\|family))* | | | | |
| | Parent temp | 2 | 1.088158 | 0.5804 |
| | Offspring temp | 2 | 1237.628 | <0.001 |
| | Parent temp:Offspring temp | 4 | 6.578507 | 0.1599 |
| *Egg hatching success (GLMM: ~ Parent temp * Offspring temp + (1\|family))* | | | | |
| | Parent temp | 2 | 4.0477 | 0.1321 |
| | Offspring temp | 2 | 64.4464 | <0.001 |
| | Parent temp:Offspring_temp | 4 | 5.7990 | 0.2147 |
| *Growth rate (LMM: ~ Parent temp * Offspring temp + Plate replicate + (1\|family))* | | | | |
| | Parent temp | 2 | 1.6445 | 0.4394 |
| | Offspring temp | 2 | 2752.7471 | 0 |
| | Plate replicate | 1 | 2.4498 | 0.1175 |
| | Parent temp:Offspring temp | 4 | 33.3274 | <0.001 |
| *Larval survival from 1st-3rd instar (LMM: ~ Parent temp * Offspring temp + Plate replicate + (1\|family))* | | | | |
| | Parent temp | 2 | 5.9049 | 0.0522 |
| | Offspring temp | 2 | 6.7801 | 0.0337 |
| | Plate replicate | 1 | 1.5563 | 0.2122 |
| | Parent temp:Offspring temp | 4 | 11.0941 | 0.0255 |
| *Larval survival from 3rd instar-diapause (LMM: ~ Parent temp * Offspring temp + Plate replicate + (1\|family))* | | | | |
| | Parent temp | 2 | 4.8609 | 0.0880 |
| | Offspring temp | 2 | 15.2739 | <0.001 |
| | Plate replicate | 1 | 0.0511 | 0.8211 |
| | Parent temp:Offspring temp | 4 | 7.0994 | 0.1307 |

**Table S3:** Model-estimated marginal means with 95% CI (obtained using the R package *emmeans*) for life-history traits in parental generation.

| Temp | Sex | Mean | SE | lower.CI | upper.CI |
|---|---|---|---|---|---|
| *Larval development time* | | | | | |
| 28 | male | 20.3909 | 0.2688 | 19.8691 | 20.9263 |
| 31 | male | 20.0953 | 0.4119 | 19.3016 | 20.9216 |
| 34 | male | 18.6069 | 0.3303 | 17.9688 | 19.2677 |
| 28 | female | 26.6924 | 0.4018 | 25.9140 | 27.4941 |
| 31 | female | 24.2290 | 0.5247 | 23.2191 | 25.2828 |
| 34 | female | 22.1402 | 0.5240 | 21.1336 | 23.1947 |
| *Pupal weight* | | | | | |
| 28 | male | 176.9385 | 2.124222 | 172.7622 | 181.1149 |
| 31 | male | 170.0771 | 3.302237 | 163.5847 | 176.5695 |
| 34 | male | 180.0573 | 2.859821 | 174.4347 | 185.6799 |
| 28 | female | 222.0457 | 2.425123 | 217.2778 | 226.8137 |
| 31 | female | 225.8298 | 3.488949 | 218.9703 | 232.6893 |
| 34 | female | 236.115 | 3.813095 | 228.6182 | 243.6118 |
| *Growth rate* | | | | | |
| 28 | male | 0.1172 | 0.0014 | 0.1145 | 0.1198 |
| 31 | male | 0.1184 | 0.0021 | 0.1142 | 0.1225 |
| 34 | male | 0.1332 | 0.0018 | 0.1296 | 0.1367 |
| 28 | female | 0.1020 | 0.0015 | 0.0990 | 0.1050 |
| 31 | female | 0.1115 | 0.0022 | 0.1071 | 0.1159 |
| 34 | female | 0.1243 | 0.0024 | 0.1195 | 0.1291 |
| *Pupal development time* | | | | | |
| 28 | male | 12.9516 | 0.0925 | 12.7709 | 13.1348 |
| 31 | male | 10.5582 | 0.1173 | 10.3301 | 10.7913 |
| 34 | male | 9.5195 | 0.0916 | 9.3412 | 9.7013 |
| 28 | female | 13.0572 | 0.1065 | 12.8494 | 13.2683 |
| 31 | female | 10.8667 | 0.1261 | 10.6217 | 11.1174 |
| 34 | female | 9.5243 | 0.1239 | 9.2838 | 9.7710 |

**Table S4:** Model-estimated marginal means/probabilities averaged over plate replicates with 95% CI (obtained using the R package *emmeans*) for life-history traits in the offspring generation.

| Parent temp | Offspring temp | Mean/Probability | SE | lower.CI | upper.CI |
|---|---|---|---|---|---|
| *Egg development time* | | | | | |
| 28 | 28 | 13.3957 | 0.1635 | 13.0750 | 13.7242 |
| 31 | 28 | 13.4089 | 0.1825 | 13.0516 | 13.7760 |
| 34 | 28 | 13.0194 | 0.1467 | 12.7314 | 13.3140 |
| 28 | 31 | 11.3184 | 0.1366 | 11.0503 | 11.5930 |
| 31 | 31 | 11.3416 | 0.1543 | 11.0393 | 11.6520 |
| 34 | 31 | 11.3540 | 0.1280 | 11.1028 | 11.6109 |
| 28 | 34 | 10.5302 | 0.1271 | 10.2807 | 10.7856 |
| 31 | 34 | 10.3401 | 0.1407 | 10.0646 | 10.6232 |
| 34 | 34 | 10.3545 | 0.1167 | 10.1254 | 10.5888 |
| *Egg hatching success* | | | | | |
| 28 | 28 | 0.7512 | 0.0481 | 0.6449 | 0.8339 |
| 31 | 28 | 0.8435 | 0.0377 | 0.7539 | 0.9046 |
| 34 | 28 | 0.8320 | 0.0332 | 0.7558 | 0.8879 |
| 28 | 31 | 0.7130 | 0.0524 | 0.5996 | 0.8047 |
| 31 | 31 | 0.8308 | 0.0401 | 0.7364 | 0.8962 |
| 34 | 31 | 0.7934 | 0.0388 | 0.7063 | 0.8598 |
| 28 | 34 | 0.6397 | 0.0589 | 0.5172 | 0.7464 |
| 31 | 34 | 0.8052 | 0.0446 | 0.7020 | 0.8788 |
| 34 | 34 | 0.7693 | 0.0419 | 0.6765 | 0.8418 |
| *Larval growth rate* | | | | | |
| 28 | 28 | 0.0554 | 0.0014 | 0.0528 | 0.0581 |
| 31 | 28 | 0.0554 | 0.0014 | 0.0526 | 0.0582 |
| 34 | 28 | 0.0571 | 0.0012 | 0.0547 | 0.0595 |
| 28 | 31 | 0.0750 | 0.0014 | 0.0724 | 0.0777 |
| 31 | 31 | 0.0732 | 0.0014 | 0.0704 | 0.0761 |
| 34 | 31 | 0.0714 | 0.0012 | 0.0691 | 0.0738 |
| 28 | 34 | 0.0834 | 0.0014 | 0.0807 | 0.0860 |
| 31 | 34 | 0.0795 | 0.0014 | 0.0766 | 0.0823 |
| 34 | 34 | 0.0795 | 0.0012 | 0.0771 | 0.0818 |
| *Larval survival (1st-3rd instar)* | | | | | |
| 28 | 28 | 0.9711 | 0.0086 | 0.9485 | 0.9840 |
| 31 | 28 | 0.9829 | 0.0065 | 0.9641 | 0.9919 |
| 34 | 28 | 0.9396 | 0.0117 | 0.9120 | 0.9589 |
| 28 | 31 | 0.9729 | 0.0080 | 0.9517 | 0.9849 |
| 31 | 31 | 0.9722 | 0.0089 | 0.9483 | 0.9853 |
| 34 | 31 | 0.9733 | 0.0071 | 0.9552 | 0.9842 |
| 28 | 34 | 0.9410 | 0.0131 | 0.9091 | 0.9621 |
| 31 | 34 | 0.9764 | 0.0078 | 0.9550 | 0.9878 |
| 34 | 34 | 0.9506 | 0.0104 | 0.9257 | 0.9675 |
| *Larval survival (3rd instar-diapause)* | | | | | |
| 28 | 28 | 0.9759 | 0.0101 | 0.9454 | 0.9895 |
| 31 | 28 | 0.9670 | 0.0123 | 0.9322 | 0.9842 |
| 34 | 28 | 0.9257 | 0.0191 | 0.8781 | 0.9556 |
| 28 | 31 | 0.9797 | 0.0086 | 0.9538 | 0.9912 |
| 31 | 31 | 0.9691 | 0.0120 | 0.9344 | 0.9857 |
| 34 | 31 | 0.9738 | 0.0087 | 0.9499 | 0.9865 |
| 28 | 34 | 0.9926 | 0.0041 | 0.9779 | 0.9976 |
| 31 | 34 | 0.9767 | 0.0096 | 0.9482 | 0.9897 |
| 34 | 34 | 0.9697 | 0.0098 | 0.9433 | 0.9841 |
